# Development and validation of LC-MS/MS bioanalytical methods to support the clinical pharmacology studies of COMP360, a form of synthetic psilocybin

**DOI:** 10.64898/2026.11.19.767836

**Authors:** Steve Silvester, Louise Wray, Helen Dooner, Gary Gilmour, Amesha Patel

## Abstract

COMP360, a form of synthetic psilocybin (PSY), is being developed for the treatment of treatment-resistant depression (TRD) and post-traumatic stress disorder (PTSD). Orally-administered PSY is rapidly cleaved to the active moiety, psilocin (PSI), which undergoes further metabolism to psilocin-O-glucuronide (PSIG) and 4-hydroxyindole-3-acetic acid (4-HIAA).

Methods for the quantification of PSY and its metabolites in plasma (PSY – 0.5 to 500 ng/mL; PSI – 0.5 to 200 ng/mL; PSIG – 1.0 to 400 ng/mL; 4-HIAA – 5 to 2000 ng/mL), and PSI in human urine (0.5 to 200 ng/mL), were developed and validated across 2 vendor bioanalytical laboratories. Plasma samples were processed by simple protein precipitation, and the extracts evaporated to dryness and reconstituted in 0.1% formic acid in water. Chromatographic separation of analytes from endogenous components, and from one another, was effected using a Waters HSS T3 analytical column and detection was performed using tandem mass spectrometry.

Method validation was conducted, and met acceptance criteria, according to ICH M10 guidance. Appropriate reference standards were sourced for each analyte and stable isotopically-labelled versions of each analyte were used as internal standards. Suitable conditions for sample handling and storage were assessed, and acceptable stability results were validated. Stability conditions were limited by the facile oxidation of psilocin but plasma samples could be kept thawed on wet ice for 8h or frozen at -80°C for 6 months, and subjected to 4 freeze/thaw cycles. The validated ranges were shown to be appropriate for the concentrations determined in samples taken in subsequent clinical studies, and data reliability was demonstrated from successful ISR analysis. Comparative performance of the multi-analyte method, across vendor laboratories, was shown through conduct of a cross-validation study employing spiked-QC and incurred samples.

**HIGHLIGHTS:**

- LC-MS/MS methods for quantification of psilocybin, psilocin and major metabolites in human plasma
- Summary of validation results for method performance and analyte stability
- Application of method in a PK study of healthy volunteers and collated ISR analysis results
- Cross-validation assessment of methods established at 2 bioanalytical vendor laboratories

## 1. Introduction

Psilocybin, 3-[2-(dimethylamino)ethyl]-1H-indol-4-yl dihydrogen phosphate (Figure 1) is one of a number of classic psychedelic compounds that are being researched for the treatment of several psychiatric conditions [1]. Psilocybin (PSY) acts as a prodrug and, once administered, it is rapidly enzymatically cleaved to the active moiety, psilocin. Psilocin (PSI) has potent agonist activity at several serotonin receptors, the most important of which is the 5-hydroxytryptamine 2A (5-HT_2A_) receptor [2,3]. This agonism is thought to be the principal driver of the human psychedelic experience and, by extension, may be significantly implicated in mediation of efficacy [4,5]. PSI subsequently undergoes further metabolism via glucuronidation to psilocin-O-glucuronide (PSIG) or oxidative deamination to 4-hydroxyindole-3-acetic acid (4-HIAA) [6,7].

**Figure 1.**
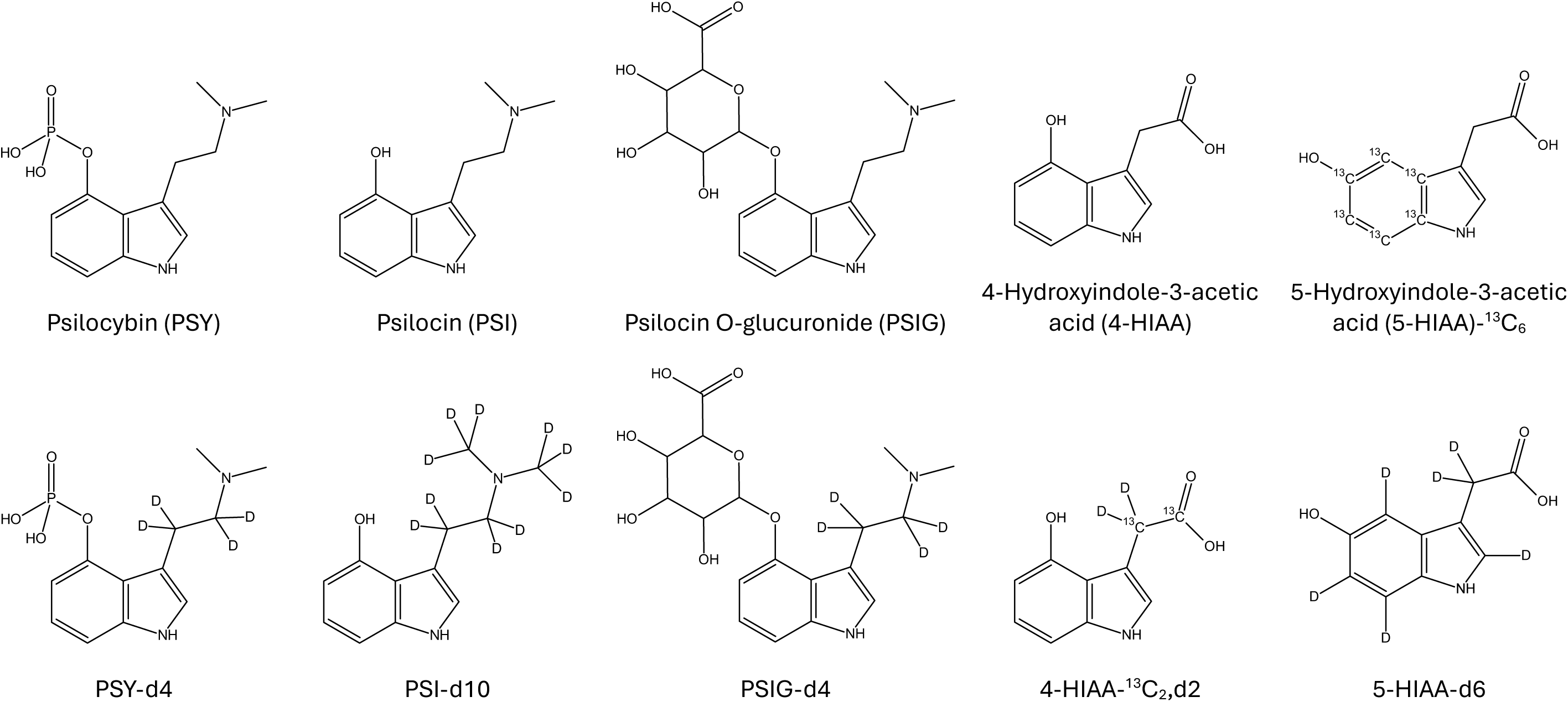
Chemical structures of psilocybin and its metabolites, and their stable isotopically-labelled internal standards

Compass is developing a form of synthetic PSY (COMP360) for the treatment of treatment-resistant depression (TRD) and post-traumatic stress disorder (PTSD) in adults. As part of clinical development, determination of pharmacokinetics (PK) following PSY (COMP360) administration is critical to understand the relationship with pharmacodynamics (PD), and to compare with exposures from preclinical toxicology studies. Consequently, a validated bioanalytical method is required for quantification of drug substance, active molecule and major metabolites in human matrices.

Determination of PSY, PSI and metabolites in biological samples is challenging. These analytes are highly polar and of low molecular weight, and there are numerous closely-related species present endogenously (e.g. tryptophan metabolites), which can give rise to interferences in detection/quantification. Historically, analytical methods for PSI have used high performance liquid chromatography (HPLC) coupled to electrochemical detection (ECD) [8–10] or gas chromatography, following off-line derivatisation, with mass spectrometry [11,12]. Increasingly, newer methods take advantage of modern LC-MS instrumentation, being highly selective and sensitive, and providing effective and efficient solutions to the challenges mentioned [13–19].

PSI and to a lesser extent PSY and 4-HIAA are known to be highly susceptible to oxidation either enzymatically in biological matrix [6,20,21] or chemically through exposure to light [22], air [10,15,20,23], transition metal ions (e.g., Fe^3+^) [23] and/or high pH [23,24]. Therefore, it is of paramount importance to the development of a successful quantitative method that satisfactory control of conditions, under which solutions and samples are prepared, handled and stored, can be maintained. Several authors have reported the improvement in stability afforded by the addition of an antioxidant, e.g., ascorbic acid (AA) or dithiothreitol (DTT), to solutions especially when highly aqueous [8,25], and/or minimising light exposure [22].

Supplementation of human plasma and urine samples with AA prior to extraction of psilocin has also been reported. For example, Hasler et al. [8] found that PSI and 4-HIAA were highly unstable in aqueous solution unless supplemented with AA and added the antioxidant to plasma samples immediately before extraction by in vitro-microdialysis. Lindenblatt et al. [10] also found that conventional off-line solid phase extraction (SPE) was unsuitable for extraction of PSI from human plasma due to extensive and variable decomposition occurring on the SPE sorbent but acceptable recovery could be achieved using a liquid-liquid extraction employing a drydown step. Martin et al [15] have also shown the effects of air and nitrogen on drydown following extraction and the protection afforded by addition of AA. More recent publications [13] have described methods that employ protein precipitation with organic solvent to extract PSI (and 4-HIAA), offering reported acceptable assay performance without the need for stabilization during extraction.

To a degree, instability during sample extraction can be mitigated once an appropriate internal standard has been added to the sample which compensates for changes in yield. However, stability of the analyte in the matrix sample from point of collection to point of analysis must be wholly maintained in order to generate a reliable quantitative result. In this regard, Kolacynska et al [13] reported acceptable stability of PSI (and 4-HIAA), according to regulatory guidance, in human plasma for 3 freeze/thaw cycles, 8h at room temperature and 1 month at -20°C, broadly in line with that shown by others [15]. Alternatively, the method reported by Brown et al [17] supplemented blood plasma (and urine) samples with AA to a concentration of 25 mM, and loss of psilocin at room temperature for 24 h or over five freeze–thaw cycles was reported to be less than 3%. In the absence of antioxidant additives, PSI appears to show slightly greater stability under frozen storage at -20°C in urine [26] and serum [16,26] than in plasma. However, it is particularly susceptible to degradation in whole blood that has been frozen, which is postulated to be as a result of red blood cell lysis, mass release of enzymes and subsequent oxidative degradation [15,19]. These differences in stability likely reflect the intrinsic antioxidant status of the biological sample itself [27,28], and effectors such as additives upon collection (e.g., blood anticoagulant [29]) and subsequent handling.

The detection of PSIG is useful in forensic toxicology applications as a surrogate marker for systemic exposure to PSY/PSI [19,30]. It is reported to be the major metabolite of PSY/PSI excreted in urine and sample handling is more straightforward, owing to its increased stability in biological matrix [19,26] (likely as a consequence of the indole-hydroxy group being conjugated and supressing susceptibility to oxidation). Several literature methods [9,11–13,31,32], in the absence of an appropriate reference standard, have quantified PSIG indirectly by determining PSI concentrations in samples before and after enzyme hydrolysis and comparing the results – the increase in concentration after deconjugation being the concentration of PSIG present.

Efficient hydrolysis of PSIG can be achieved by using β-glucuronidase but the extent of conversion and the degradation of PSI are dependent on the incubation conditions (pH, temperature and time). Addition of AA to incubations has been shown to be protective when instability has been observed (primarily in serum samples) and optimised methodologies have been reported to deliver complete conversion of PSIG and protect against losses of PSI in human serum and urine [26,31,32].

For absolute quantification of PSIG, this indirect approach is non-ideal. Aside from the added complexity of the workflow, and the intrinsic increase in error arising from 2 separate analyses, the determination is also reliant on both complete conversion of PSIG and insignificant degradation of PSI during the hydrolysis procedure for every sample analysed. The lack of a reference standard presents a challenge in ensuring quality control of each batch. Moreover, in order to comply with regulatory bioanalytical method validation guidelines, a reference standard, which is well characterised, of appropriate quality (e.g., purity, identity) and obtained from an authentic and traceable source, is an absolute requirement.

The aim of the work undertaken was to develop and validate a single bioanalytical method for the absolute determination of drug substance, PSY, the active metabolite PSI, and the 2 major metabolites PSIG and 4-HIAA in human plasma to provide accurate and reliable concentration data in clinical pharmacology studies sponsored by Compass. Due to challenges experienced within method development, and described herein, PSY was omitted from the multi-analyte assay and a separate method developed and validated. A single-analyte method for the determination of PSI in human urine was also generated. Validation data from all methods are reported from 2 vendor bioanalytical laboratories, along with results from a cross-validation study to compare performance of the multi-analyte plasma method at both. Additionally, in-study application of the method is described to include representative concentration-time data in healthy human participants and incurred sample reproducibility (ISR) analysis results spanning 5 clinical trials.

## 2. Experimental

### 2.1 Reference compounds, chemicals and reagents

Reference compounds were procured from various sources. PSY, as COMP360, was manufactured at Onyx Scientific (Sunderland, UK). PSI was used as manufactured material from Onyx Scientific or purchased as certified solutions (1 mg/mL, Supelco from Merck (Gillingham, UK). 4-HIAA and PSIG were manufactured at Onyx Scientific and Apex Molecular (Macclesfield, UK), respectively; PSIG was also obtained from Alsachim (Strasbourg, France). Stable isotopically-labelled Internal standards were employed in the work. PSY-d4, PSIG-d4 and 4-HIAA-^13^C_2_,d2 were manufactured at Apex Molecular. 5-HIAA-^13^C_6_ (5-hydroxyindoleacetic acid) was purchased as solid material from Merck and PSI-d10 and 5-HIAA-d6 were purchased as certified solutions (Supelco and Cerilliant, respectively) from Merck.

Acetonitrile (MeCN) and methanol (MeOH), of HPLC-grade or higher, were purchased from Fisher Scientific (Loughborough, UK) or VWR (Lutterworth, UK); ultrapure water was prepared in-house (Milli-Q or Elga systems). Formic acid (FA, AR-grade) and L-ascorbic acid (AA, >99%) were purchased from either Merck (Gillingham, UK) or Fisher.

Blank human blood/plasma, collected in K2EDTA, was procured from BioIVT (West Sussex, UK). Blank human urine was collected in house from healthy volunteers.

### 2.2 Preparation of stock solutions

Primary stock solutions of PSY, PSI, PSIG and 4-HIAA were prepared at a nominal concentration of 1mg/mL by dissolving solid material in appropriate amounts of water containing 25 mM AA. Solutions were prepared in amber (or foil-wrapped) glass vials, typically sub-aliquoted for single use and stored at -80°C. Thawed solutions were kept on wet ice during use. For PSI, certified reference standard prepared at a concentration of 1 mg/mL in MeCN was also used.

Stock solutions of SIL-IS compounds were prepared at nominal concentrations of either 1 or 0.1 mg/mL by dissolving solid material in the same solvents used for the unlabelled compounds, and handled similarly. Certified reference standards prepared at a concentration of 0.1 mg/mL were also used for PSI-d10 (in MeCN) and 5-HIAA-d6 (in MeOH).

Stock solutions of PSY or PSIG/4-HIAA were stable in water containing 25 mM AA for at least 52 or 146 days, respectively, at -80°C. Stock solutions of PSI were stable in MeCN or in water containing 25 mM AA for at least 146 or 117 days, respectively, at -80°C.

### 2.3 Preparation of calibration and quality control samples

Calibration samples were prepared by fortifying blank matrix with each analyte at a minimum of 8 concentration levels across the calibration range (see Table 1). Quality Control (QC) samples were prepared similarly at a minimum of lower limit of quantification (LLOQ), low, medium, high and upper limit of quantification (ULOQ) levels (corresponding to concentrations at the assay LLOQ, approximately three times the LLOQ, 30 – 50% of the ULOQ, 80% of the ULOQ and the ULOQ, respectively). Dilution QC samples were also prepared at a concentration above the calibration range (6 to 20-fold). Both sets of samples were prepared by spiking primary stock solutions directly into blank matrix, and performing serial dilutions in matrix to the required concentrations, or by preparing a set of working solutions from the primary stock solutions and spiking an aliquot of each into volumes of blank matrix (at a minimum dilution factor of 20-fold). In the case of the latter, working solutions were prepared in comparable solvents used to prepare primary solutions [i.e.,25 mM AA in water or 90:10 (v/v) water/MeCN] and were stored under the same conditions (typically as single-use aliquots). Separate preparations of primary stock solutions of each analyte were used for the preparation of calibration and quality control samples.

**Table 1.** Summary of defining parameters for validated bioanalytical methods.

| ID | Method | Matrix | Sample volume (µL) | Method details | Analyte calibration concentration range (ng/mL) |  |  |  |
|---|---|---|---|---|---|---|---|---|
|  |  |  |  |  | PSY | PSI | PSIG | 4-HIAA |
| 1 | Single-analyte<br>CRO-A | Human plasma (EDTA) | 50 | Extraction – A<br>Instrumentation- A<br>Column - A | 0.500 - 500 | N/A | N/A | N/A |
| 2 | Multi-analyte<br>CRO-A | Human plasma (EDTA) | 50 | Extraction – B<br>Instrumentation- A<br>Column -B | N/A | 0.500 - 200 | 0.892 – 357<sup>a</sup> | 5.00 - 2000 |
| 3 | Multi-analyte<br>CRO-B | Human plasma (EDTA) | 50 | Extraction – C<br>Instrumentation- B<br>Column -B | N/A | 0.500 - 200 | 1.00 - 400 | 5.00 - 2000 |
| 4 | Single analyte<br>CRO-B | Human urine | 25 | Extraction – D<br>Instrumentation- B<br>Column -B | N/A | 0.500 - 200 | N/A | N/A |

Extraction: A) Extraction using 0.1% FA in MeCN, supernatant drydown/reconstitute ; B) extraction using MeCN on Ostro plate, eluate drydown/reconstitute; C) Extraction using MeCN, supernatant drydown/reconstitute ; D) Dilution with 25mM AA in 90:10 (v/v) water/MeCN
Instrumentation: A) ExionLC AD, API6500+; B) Acquity I-Class (FTN), Xevo TQ-XS
Column: A) Acquity HSS T3 2.1 x 50 mm, 1.8 µm; B) Acquity HSS T3 2.1 x 100 mm, 1.8 µm
a Range amended from nominal to correct for salt form content

### 2.4 Sample extraction

Sample extraction protocols differed between methods and vendor laboratories (see Table 1). Plasma methods were based on protein precipitation utilizing MeCN as extraction solvent and the urine method employed sample dilution. Calibration, QC and test samples were thawed and kept on wet ice prior to extraction. Sampling of aliquots into polypropylene tubes or 96-well plates was also performed on wet ice until extraction into MeCN (or dilution in urine method) had been effected.

In plasma methods, an aliquot (20 μL) of aqueous solution containing the relevant SIL-IS internal standards for each analyte (diluted from primary stocks to appropriate concentrations) was added to sample aliquots (50 µL) in polypropylene tubes or 96-well plate and mixed thoroughly to ensure homogeneity. Subsequently, an excess (200 µL) of MeCN was added to each sample and mixed thoroughly. Following centrifugation (at 13000g or greater), supernatants were subjected to drydown under a stream of nitrogen (30 or 40°C) and subsequently reconstituted in a volume of water containing 0.1% FA (100 or 150 µL). In the human metabolites method validated at CRO-A (method 2 in Table 1), protein precipitation of the sample was performed in Ostro Lipid Removal 96-well plates (25 mg, Waters, Milford, MA). Extracted samples were kept in sealed 96-well plates refrigerated at ≤10°C.

For the urine method, an aliquot (500 μL) of 25mM AA in 90:10 water/MeCN, containing PSI-d10 internal standard (diluted from primary stock), was added to sample aliquots (25 µL) and mixed thoroughly. Following centrifugation (at 16100g), supernatants (150 µL) were recovered and sealed 96-well plates refrigerated at ≤10°C.

### 2.5 LC-MS/MS instrumentation and method

Analytes in sample extracts were chromatographically separated and detected/quantified by tandem mass spectrometry with electrospray ionization. Instrumentation differed between the vendor laboratories as summarized in Table 1. Chromatography was performed on an Acquity I-Class (FTN) (Waters, Milford, MA) or Sciex ExionLC AD (Sciex, Marlborough, MA) system. Mass spectrometric detection was performed on a Sciex API6500+ or Waters Xevo TQ-XS. Separation was achieved on a Waters Acquity HSS T3 (2.1 x 100 mm or 2.1 x 50 mm, 1.8 μm) column, maintained at 50°C. Gradient elution was employed with the aqueous phase modified with 0.1% FA and an organic phase of 0.1:100 (v/v) FA/MeCN or simply MeCN. The flow rate ranged from 0.5 to 0.6 mL/min. Specific details are provided in Table 2.

**Table 2.** Gradient elution conditions employed in validated bioanalytical methods.

| Step | Method 1 |  |  | Method 2/3 |  |  | Method 4 |  |  |
|---|---|---|---|---|---|---|---|---|---|
|  | Time (mins) | %A | %B | Time (mins) | %A | %B | Time (mins) | %A | %B |
| 1 | 0.00 | 97 | 3 | 0.00 | 98 | 2 | 0.00 | 98 | 2 |
| 2 | 1.20 | 90 | 10 | 0.20 | 98 | 2 | 0.20 | 98 | 2 |
| 3 | 1.30 | 10 | 90 | 3.20 | 75 | 25 | 2.50 | 80 | 20 |
| 4 | 1.80 | 10 | 90 | 3.30 | 5 | 95 | 3.00 | 5 | 95 |
| 5 | 1.90 | 97 | 3 | 3.90 | 5 | 95 | 4.00 | 5 | 95 |
| 6 | 2.40 | 97 | 3 | 4.10 | 98 | 2 | 4.10 | 98 | 2 |
| 7 | N/A | N/A | N/A | 5.00/7.00 | 98 | 2 | 6.00 | 98 | 2 |

1) A - 0.1% FA in water; B – 0.1% FA in MeCN; 0.6 ml/min, 50°C, switch from waste to MS at 0.6 min
2) A - 0.1% FA in water; B – MeCN; 0.55 ml/min, 50°C, switch from waste to MS at 0.7 min and back to waste at 4 min; end of re-equilibration at 5 min
3) A - 0.1% FA in water; B – 0.1% FA in MeCN; 0.5 ml/min, 50°C, switch from waste to MS at 1.2 min and back to waste at 4 min; end of re-equilibration at 7 min
4) A - 0.1% FA in water; B – 0.1% FA in MeCN; 0.5 ml/min, 50°C, switch from waste to MS at 1.6 min and back to waste at 2.6 min

PSY, PSI and PSIG were ionized under positive electrospray ionization and 4-HIAA under negative ionization. Standard source parameters and details of the MRM (multiple reaction monitoring) experiments utilized are summarized in Table 3. Analyst v1.7 (with Multiquant v3.0.3) or MassLynx v4.2 (with TargetLynx v4.1) software was used to operate the Sciex or Waters LC-MS/MS system and process data, respectively.

**Table 3.** Summary of mass spectrometry parameters for psilocybin and its metabolites, and their stable isotopically-labelled internal standards.

| Analyte | Q1/Q3 mass (amu)<sup>a</sup> | Collision energy (eV)<sup>b</sup> | Method employed |
|---|---|---|---|
| PSY | 285/205 | 25 | 1 |
| PSY-d4 | 289/209 | 25 | 1 |
| PSI | 205/160 | 25 | 2 |
| PSI | 205/58 | 15 | 3, 4 |
| PSI-d10 | 215/164 | 25 | 2 |
| PSI-d10 | 215/66 | 15 | 3, 4 |
| PSIG | 381/205 | 20 | 2, 3 |
| PSIG-d4 | 385/209 | 20 | 2, 3 |
| 4-HIAA | 190/131 | -35 | 2 |
| 4-HIAA | 190/146 | -15 | 3 |
| 5-HIAA-<sup>13</sup>C<sub>6</sub> | 196/150 | -30 | 2 |
| 5-HIAA-d6 | 196/152 | -15 | 3 |
| 4-HIAA-<sup>13</sup>C<sub>2</sub>,d2 | 194/149 | -15 | 3 |

Standard instrument parameters for API6500:  IonSpray voltage, 5500/-4500 V; Temp, 600°C; Gas 1, 60 psi; Gas 2, 60 psi; Curtain gas, 40 psi; Collision gas, 8; Declustering potential, 50/-60 V
Standard instrument parameters for Xevo TQ-XS:  Capillary voltage, 3.5/-2.5 kV; Desolvation temp, 500°C; Source temp, 150°C; Desolvation gas flow, 550 L/h; Cone gas flow, 150 L/h; Collision gas flow, 0.20 mL/min; Cone voltage, 30/-30 V
a	Nominal masses presented (amended to optimize sensitivity)
b	Typical value presented (amended to optimize sensitivity)

### 2.6 Method validation

A summary of the method validations performed is given in Table 1. The studies were all conducted in GLP-compliant laboratories and the experiments performed were in accordance with the requirements laid out in the current International Council for Harmonisation (ICH) M10 guidance [33].

#### 2.6.1 Calibration curve and range

The relationship between nominal analyte concentration and response, in each method, was assessed using calibration standards in analytical runs. The detector response ratio for analyte:internal standard was plotted against nominal concentration and the simplest regression model that adequately described the relationship applied.

The back-calculated concentration for each analyte, for each standard, needed to be within ±15% of the nominal concentration (± 20% at the LLOQ). Standards that did not meet these criteria were excluded from the regression analysis. At least 75% (and a minimum of 6) of the calibration standards needed to meet these criteria. Calibration curves were freshly prepared for each validation run containing stability samples and in other runs until frozen stability had been demonstrated.

#### 2.6.2 Accuracy and precision including dilution integrity

The intra-(within) and inter-(between) batch assay accuracy and precision were determined by the analysis of replicates (n = 6) of QC samples, spanning the method range, along with calibration standards. A minimum of 3 analytical runs were performed with at least one employing freshly prepared calibration standards and/or QC samples and one of a size equivalent to that of a prospective analytical run of study samples.

Dilution integrity was assessed by analyzing the dilution QC sample following appropriate dilution into the calibration range (n = 6) in a single batch.

#### 2.6.3 Selectivity and specificity

Assay selectivity, with respect to potential interference arising in blank matrix, was assessed by analysing at least six different lots of blank matrix for the response of each analyte and internal standard. A haemolysed source (≥2% haemolysis) and a lipemic source were also evaluated for plasma assays.

Specificity, with respect to potential interference arising between analytes and/or internal standard was assessed by analysing a single source of blank matrix individually-spiked with each analyte, at the ULOQ concentration, or internal standard. Additional specificity testing, with respect to potential concomitant medications, was also performed through a paper-based assessment of molecular weight/lipophilicity or analysis of blank matrix and low/high QC samples spiked with compounds at pharmacologically-relevant concentrations.

#### 2.6.4 Matrix effect/recovery

The potential effect of the matrix on each analyte (through ionization suppression/enhancement and/or recovery) was evaluated in each method validation by determining the matrix factor (in different lots of matrix) and recovery separately, or the overall matrix effect (in different lots of plasma). The effect of haemolysed matrix, and lipemic matrix in human methods, was also assessed.

To determine the matrix factor, different sources of blank matrix (at least 6), spiked post-extraction with each analyte at nominal low and high QC concentrations, were analysed and the peak responses compared, with and without normalization of internal standard, against those determined from spiked solutions in the absence of matrix. Recovery was determined in a single source of matrix by spiking each analyte before or after extraction, at low, mid and high QC concentrations, and comparing the resultant peak responses.

The overall matrix effect was assessed by spiking different sources of blank matrix (at least 6) with each analyte at nominal low and high QC concentrations (≥ 3 replicates) and determining the concentrations in a run against calibration samples.

#### 2.6.5 Stability

The stability of analytes in matrix, under conditions reflective of sample collection, storage and processing, were investigated at low and high QC sample concentrations. Sample replicates (n = 6) were analysed following storage frozen at -20 and/or -80°C, storage thawed at room temperature or on wet ice, and multiple cycles of freezing and thawing. Stability (and non-specific adsorption) was also assessed for PSI in urine samples under conditions representative of collection/storage within the clinic (amber polyethylene containers kept at approximately 4°C). Stability samples were analysed in batches using freshly-prepared calibration samples and included a standard set of QC samples for purposes of batch acceptance. Stability in whole blood (samples spiked at 37°C and then plunged in wet ice) and in extracts at ≤10°C, following sample processing, was also assessed. Details of the storage durations and freeze/thaw cycles tested are given in Table 5.

**Table 4.**
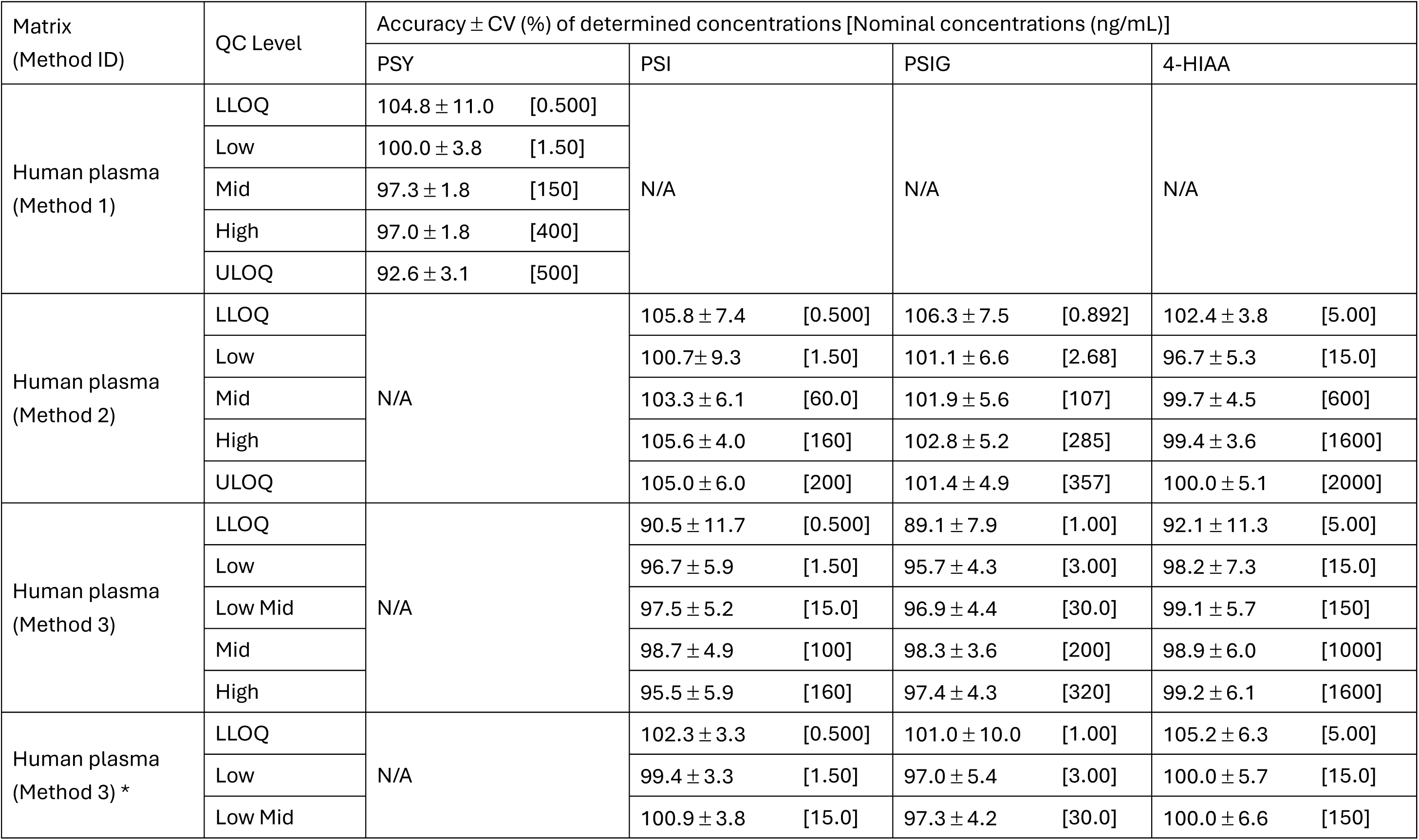
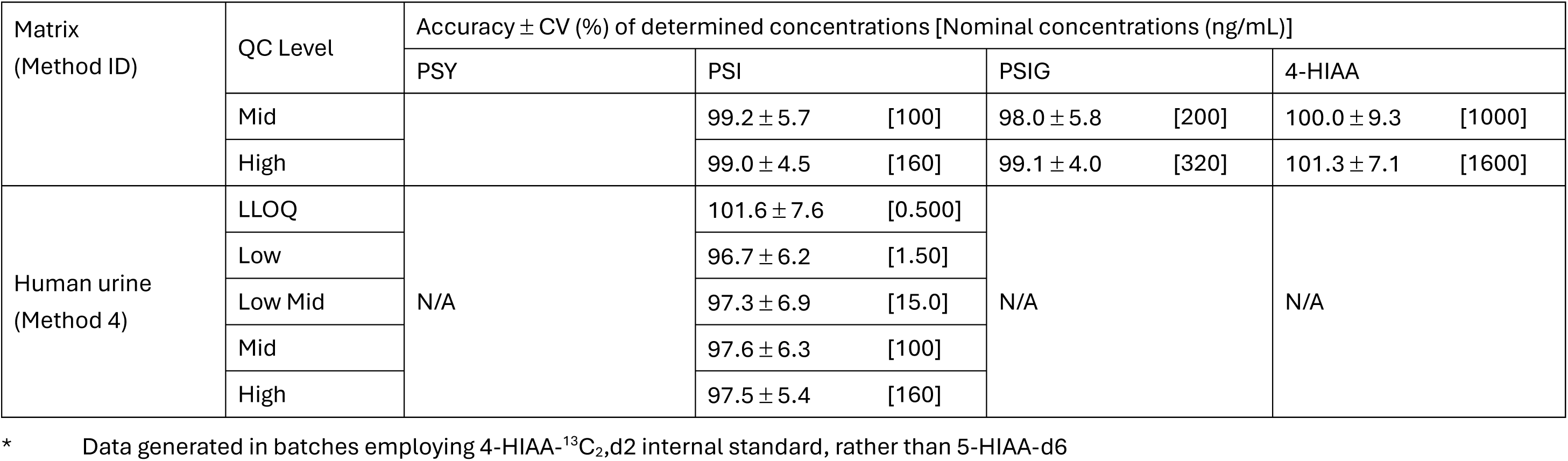
Inter-batch accuracy and precision data determined in QC samples.

| Matrix<br>(Method ID) | QC Level | Accuracy ± CV (%) of determined concentrations [Nominal concentrations (ng/mL)] |  |  |  |
|---|---|---|---|---|---|
|  |  | PSY | PSI | PSIG | 4-HIAA |
| Human plasma<br>(Method 1) | LLOQ | 104.8 ± 11.0 [0.500] | N/A | N/A | N/A |
|  | Low | 100.0 ± 3.8 [1.50] |  |  |  |
|  | Mid | 97.3 ± 1.8 [150] |  |  |  |
|  | High | 97.0 ± 1.8 [400] |  |  |  |
|  | ULOQ | 92.6 ± 3.1 [500] |  |  |  |
| Human plasma<br>(Method 2) | LLOQ | N/A | 105.8 ± 7.4 [0.500] | 106.3 ± 7.5 [0.892] | 102.4 ± 3.8 [5.00] |
|  | Low |  | 100.7± 9.3 [1.50] | 101.1 ± 6.6 [2.68] | 96.7 ± 5.3 [15.0] |
|  | Mid |  | 103.3 ± 6.1 [60.0] | 101.9 ± 5.6 [107] | 99.7 ± 4.5 [600] |
|  | High |  | 105.6 ± 4.0 [160] | 102.8 ± 5.2 [285] | 99.4 ± 3.6 [1600] |
|  | ULOQ |  | 105.0 ± 6.0 [200] | 101.4 ± 4.9 [357] | 100.0 ± 5.1 [2000] |
| Human plasma<br>(Method 3) | LLOQ | N/A | 90.5 ± 11.7 [0.500] | 89.1 ± 7.9 [1.00] | 92.1 ± 11.3 [5.00] |
|  | Low |  | 96.7 ± 5.9 [1.50] | 95.7 ± 4.3 [3.00] | 98.2 ± 7.3 [15.0] |
|  | Low Mid |  | 97.5 ± 5.2 [15.0] | 96.9 ± 4.4 [30.0] | 99.1 ± 5.7 [150] |
|  | Mid |  | 98.7 ± 4.9 [100] | 98.3 ± 3.6 [200] | 98.9 ± 6.0 [1000] |
|  | High |  | 95.5 ± 5.9 [160] | 97.4 ± 4.3 [320] | 99.2 ± 6.1 [1600] |
| Human plasma<br>(Method 3) * | LLOQ | N/A | 102.3 ± 3.3 [0.500] | 101.0 ± 10.0 [1.00] | 105.2 ± 6.3 [5.00] |
|  | Low |  | 99.4 ± 3.3 [1.50] | 97.0 ± 5.4 [3.00] | 100.0 ± 5.7 [15.0] |
|  | Low Mid |  | 100.9 ± 3.8 [15.0] | 97.3 ± 4.2 [30.0] | 100.0 ± 6.6 [150] |
| Matrix<br>(Method ID) | QC Level | Accuracy ± CV (%) of determined concentrations [Nominal concentrations (ng/mL)] |  |  |  |
|  |  | PSY | PSI | PSIG | 4-HIAA |
|  | Mid |  | 99.2 ± 5.7 [100] | 98.0 ± 5.8 [200] | 100.0 ± 9.3 [1000] |
|  | High |  | 99.0 ± 4.5 [160] | 99.1 ± 4.0 [320] | 101.3 ± 7.1 [1600] |
| Human urine<br>(Method 4) | LLOQ | N/A | 101.6 ± 7.6 [0.500] | N/A | N/A |
|  | Low |  | 96.7 ± 6.2 [1.50] |  |  |
|  | Low Mid |  | 97.3 ± 6.9 [15.0] |  |  |
|  | Mid |  | 97.6 ± 6.3 [100] |  |  |
|  | High |  | 97.5 ± 5.4 [160] |  |  |

\* Data generated in batches employing 4-HIAA-<sup>13</sup>C<sub>2</sub>,d2 internal standard, rather than 5-HIAA-d6

**Table 5:**
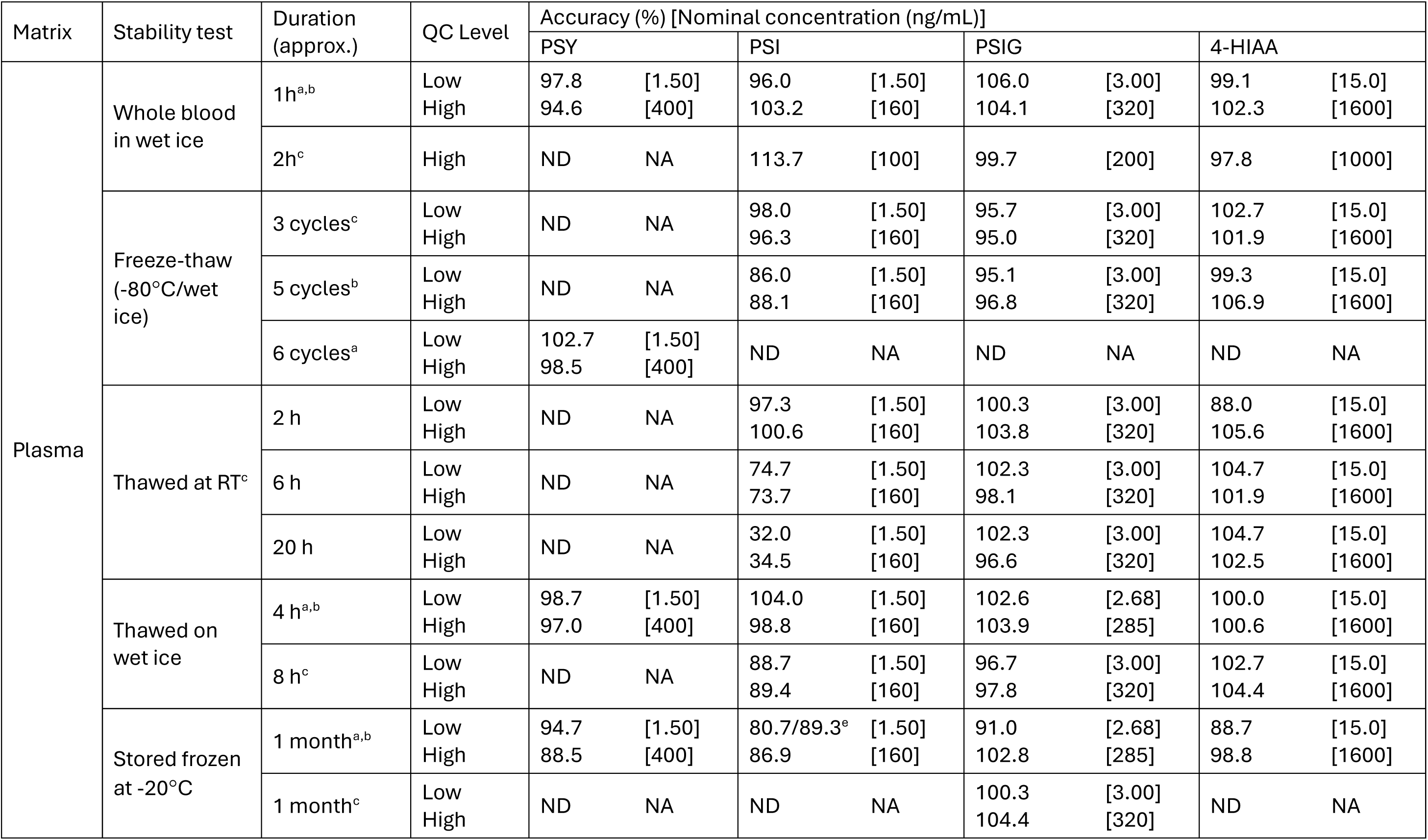
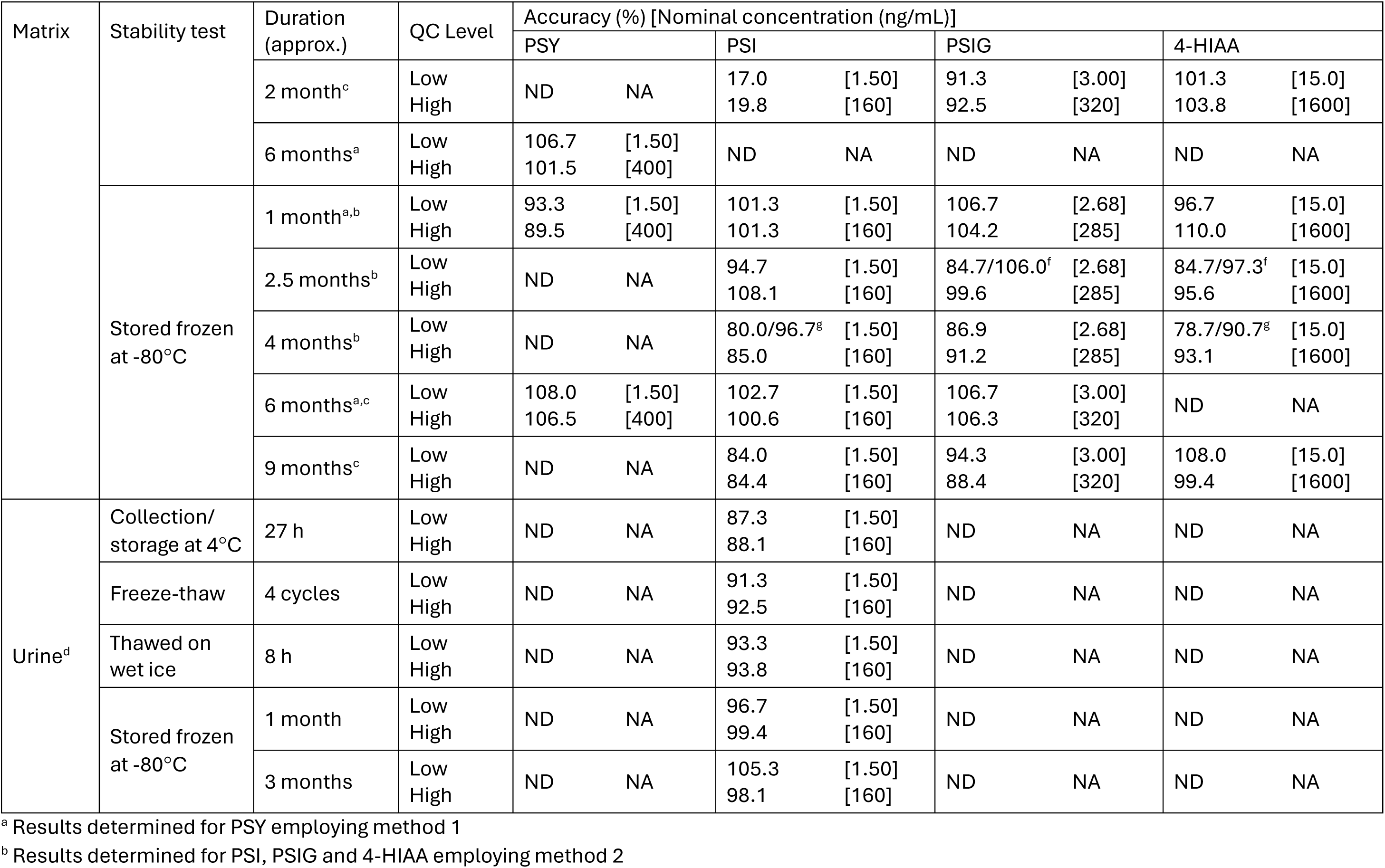
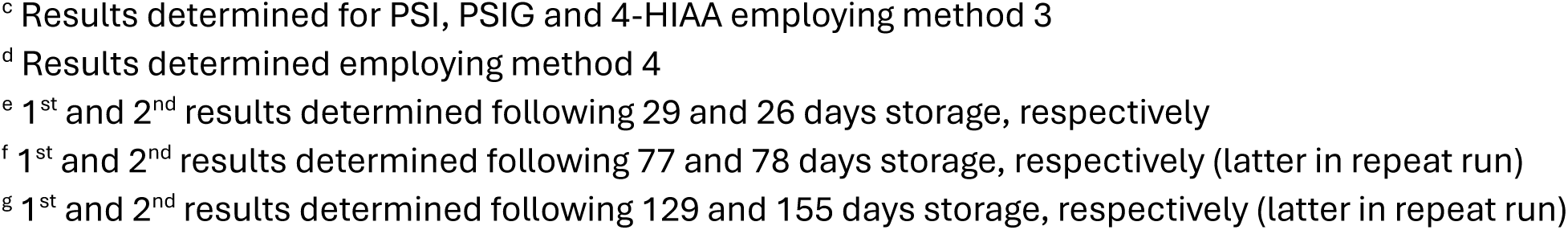
Stability results from QC samples following benchtop (thawed) and frozen storage, and freeze-thaw cycles.

| Matrix | Stability test | Duration (approx.) | QC Level | Accuracy (%) [Nominal concentration (ng/mL)] |  |  |  |  |  |  |  |
|  |  |  |  | PSY |  | PSI |  | PSIG |  | 4-HIAA |  |
|---|---|---|---|---|---|---|---|---|---|---|---|
| Plasma | Whole blood in wet ice | 1h<sup>a,b</sup> | Low<br>High | 97.8<br>94.6 | [1.50]<br>[400] | 96.0<br>103.2 | [1.50]<br>[160] | 106.0<br>104.1 | [3.00]<br>[320] | 99.1<br>102.3 | [15.0]<br>[1600] |
|  |  | 2h<sup>c</sup> | High | ND | NA | 113.7 | [100] | 99.7 | [200] | 97.8 | [1000] |
|  | Freeze-thaw (-80°C/wet ice) | 3 cycles<sup>c</sup> | Low<br>High | ND | NA | 98.0<br>96.3 | [1.50]<br>[160] | 95.7<br>95.0 | [3.00]<br>[320] | 102.7<br>101.9 | [15.0]<br>[1600] |
|  |  | 5 cycles<sup>b</sup> | Low<br>High | ND | NA | 86.0<br>88.1 | [1.50]<br>[160] | 95.1<br>96.8 | [3.00]<br>[320] | 99.3<br>106.9 | [15.0]<br>[1600] |
|  |  | 6 cycles<sup>a</sup> | Low<br>High | 102.7<br>98.5 | [1.50]<br>[400] | ND | NA | ND | NA | ND | NA |
|  | Thawed at RT<sup>c</sup> | 2 h | Low<br>High | ND | NA | 97.3<br>100.6 | [1.50]<br>[160] | 100.3<br>103.8 | [3.00]<br>[320] | 88.0<br>105.6 | [15.0]<br>[1600] |
|  |  | 6 h | Low<br>High | ND | NA | 74.7<br>73.7 | [1.50]<br>[160] | 102.3<br>98.1 | [3.00]<br>[320] | 104.7<br>101.9 | [15.0]<br>[1600] |
|  |  | 20 h | Low<br>High | ND | NA | 32.0<br>34.5 | [1.50]<br>[160] | 102.3<br>96.6 | [3.00]<br>[320] | 104.7<br>102.5 | [15.0]<br>[1600] |
|  | Thawed on wet ice | 4 h<sup>a,b</sup> | Low<br>High | 98.7<br>97.0 | [1.50]<br>[400] | 104.0<br>98.8 | [1.50]<br>[160] | 102.6<br>103.9 | [2.68]<br>[285] | 100.0<br>100.6 | [15.0]<br>[1600] |
|  |  | 8 h<sup>c</sup> | Low<br>High | ND | NA | 88.7<br>89.4 | [1.50]<br>[160] | 96.7<br>97.8 | [3.00]<br>[320] | 102.7<br>104.4 | [15.0]<br>[1600] |
|  | Stored frozen at -20°C | 1 month<sup>a,b</sup> | Low<br>High | 94.7<br>88.5 | [1.50]<br>[400] | 80.7/89.3<sup>e</sup><br>86.9 | [1.50]<br>[160] | 91.0<br>102.8 | [2.68]<br>[285] | 88.7<br>98.8 | [15.0]<br>[1600] |
|  |  | 1 month<sup>c</sup> | Low<br>High | ND | NA | ND | NA | 100.3<br>104.4 | [3.00]<br>[320] | ND | NA |
| Matrix | Stability test | Duration (approx.) | QC Level | Accuracy (%) [Nominal concentration (ng/mL)] |  |  |  |  |  |  |  |
|  |  |  |  | PSY |  | PSI |  | PSIG |  | 4-HIAA |  |
|  |  | 2 month<sup>c</sup> | Low<br>High | ND | NA | 17.0<br>19.8 | [1.50]<br>[160] | 91.3<br>92.5 | [3.00]<br>[320] | 101.3<br>103.8 | [15.0]<br>[1600] |
|  |  | 6 months<sup>a</sup> | Low<br>High | 106.7<br>101.5 | [1.50]<br>[400] | ND | NA | ND | NA | ND | NA |
|  | Stored frozen at -80°C | 1 month<sup>a,b</sup> | Low<br>High | 93.3<br>89.5 | [1.50]<br>[400] | 101.3<br>101.3 | [1.50]<br>[160] | 106.7<br>104.2 | [2.68]<br>[285] | 96.7<br>110.0 | [15.0]<br>[1600] |
|  |  | 2.5 months<sup>b</sup> | Low<br>High | ND | NA | 94.7<br>108.1 | [1.50]<br>[160] | 84.7/106.0<sup>f</sup><br>99.6 | [2.68]<br>[285] | 84.7/97.3<sup>f</sup><br>95.6 | [15.0]<br>[1600] |
|  |  | 4 months<sup>b</sup> | Low<br>High | ND | NA | 80.0/96.7<sup>g</sup><br>85.0 | [1.50]<br>[160] | 86.9<br>91.2 | [2.68]<br>[285] | 78.7/90.7<sup>g</sup><br>93.1 | [15.0]<br>[1600] |
|  |  | 6 months<sup>a,c</sup> | Low<br>High | 108.0<br>106.5 | [1.50]<br>[400] | 102.7<br>100.6 | [1.50]<br>[160] | 106.7<br>106.3 | [3.00]<br>[320] | ND | NA |
|  |  | 9 months<sup>c</sup> | Low<br>High | ND | NA | 84.0<br>84.4 | [1.50]<br>[160] | 94.3<br>88.4 | [3.00]<br>[320] | 108.0<br>99.4 | [15.0]<br>[1600] |
| Urine<sup>d</sup> | Collection/ storage at 4°C | 27 h | Low<br>High | ND | NA | 87.3<br>88.1 | [1.50]<br>[160] | ND | NA | ND | NA |
|  | Freeze-thaw | 4 cycles | Low<br>High | ND | NA | 91.3<br>92.5 | [1.50]<br>[160] | ND | NA | ND | NA |
|  | Thawed on wet ice | 8 h | Low<br>High | ND | NA | 93.3<br>93.8 | [1.50]<br>[160] | ND | NA | ND | NA |
|  | Stored frozen at -80°C | 1 month | Low<br>High | ND | NA | 96.7<br>99.4 | [1.50]<br>[160] | ND | NA | ND | NA |
|  |  | 3 months | Low<br>High | ND | NA | 105.3<br>98.1 | [1.50]<br>[160] | ND | NA | ND | NA |

<sup>a</sup> Results determined for PSY employing method 1
<sup>b</sup> Results determined for PSI, PSIG and 4-HIAA employing method 2
<sup>c</sup> Results determined for PSI, PSIG and 4-HIAA employing method 3
<sup>d</sup> Results determined employing method 4
<sup>e</sup> 1<sup>st</sup> and 2<sup>nd</sup> results determined following 29 and 26 days storage, respectively
<sup>f</sup> 1<sup>st</sup> and 2<sup>nd</sup> results determined following 77 and 78 days storage, respectively (latter in repeat run)
<sup>g</sup> 1<sup>st</sup> and 2<sup>nd</sup> results determined following 129 and 155 days storage, respectively (latter in repeat run)

### 2.7 Method application and ISR

The methods were developed and validated to support clinical pharmacology studies in the development of COMP360. To date, plasma sample analysis has been conducted in 6 clinical studies and incurred sample reproducibility (ISR) analysis was performed in 5 of these. Urine sample analysis, and assessment of ISR, has been performed in a single study. Individual study samples were selected for ISR across the concentration-time profiles for each analyte with the number analysed being a proportion of the total number of study samples (10% up to 1000 samples, plus 5% thereafter). ISR samples were analysed in a separate batch and on a separate day to first analysis. The % difference in concentration between initial and repeat analysis was calculated as follows:

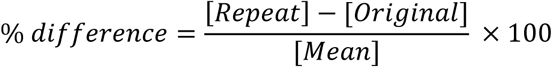

Representative data included in this article were generated from a double-blind, randomised, placebo-controlled study in which healthy volunteers received either a single dose of 25 mg COMP360 (n = 6) or placebo (n = 2) under fasting conditions.

The study was conducted in accordance with the moral, ethical and scientific principles governing clinical research as set out in the Declaration of Helsinki, and with the principles of Good Clinical Practice (GCP). Informed consent was obtained from all participants enrolled in the study.

Blood samples were collected from participants in K2EDTA coated tubes prior to dosing and at 0.25, 0.5, 0.75, 1, 1.5, 2, 2.5, 3, 4, 6, 8, 12 and 24 h post-dose. Tubes were gently inverted and centrifuged for 10 mins at 1200g and 4°C to generate plasma. Two aliquots of plasma were transferred to separate polypropylene tubes and immediately frozen at -80°C until analysis.

### 2.8 Human plasma method cross-validation

Cross-validation of the methods for the determination of PSI, PSIG and 4-HIAA in human plasma (methods 2 and 3) was performed at the respective laboratories using QC samples and incurred study samples. A batch of QC samples was prepared at CRO-B, randomized and split into 2 equivalents sets (n = 6), at each concentration level. Both sets were frozen immediately with one set retained at CRO-B and the other set dispatched to CRO-A. Similarly, 30 clinical study samples (unused backup aliquots) from a completed trial were selected, thawed and split into 2 aliquots. Both sets of aliquots were frozen immediately with one set retained at CRO-B and the other set dispatched to CRO-A. The QC samples and study samples were analysed at both laboratories using their validated methods and in batches that included an additional set of batch acceptance QC samples. All samples were analysed within validated stability limits.

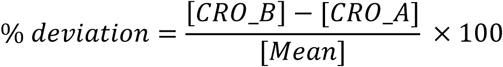

## 3. Results and discussion

### 3.1 Method development

Two separate vendor bioanalytical laboratories have been employed in the clinical development of COMP360. Aspects of method development have been performed at both, with learnings shared upon transfer of the plasma methodology from CRO-A to B. Due to differences in laboratory equipment/instrumentation and standard approaches, the final validated methods for PSI, PSIG and 4-HIAA in plasma were not absolutely uniform in all aspects but delivered the same method requirements. Target limits of quantification for each analyte were based on literature exposure information, where available. The actual limits of quantification achievable were dependent on the inherent chemistry of the analytes, matrix interferences and the need to balance optimization of each analyte in targeting development of a multi-analyte method.

The inherent instability of PSI and, to a lesser extent, PSY and 4-HIAA, is well known, as described in Section 1, and considerable attention was afforded to determining conditions under which analyte stability could be maintained in the handling and storage of solutions and samples. Stock solutions of solid compounds, and working solutions, were prepared in water containing 25mM AA and in amber (or foil-wrapped) glass vials to protect against oxidation. The solutions were typically split into smaller aliquots for single-use, stored frozen at -80°C and kept on ice when thawed. Certified reference solutions of PSI, at 1 mg/mL in MeCN, were handled similarly and were found to be acceptably stable without addition of antioxidant.

MS/MS optimization experiments yielded comparable results between laboratories and instruments with PSY, PSI and PSIG yielding highest sensitivity for the protonated molecular ion under positive ionization and 4-HIAA for the deprotonated molecular ion under negative ionization. Product ions giving highest sensitivity differed between the Sciex API6500+ and Waters Xevo TQ-XS instruments; the optimized MRM experiments developed are listed in Table 3.

Chromatography was initially investigated on a biphenyl phase with the aim of benefiting from its enhanced retention and selectivity for aromatic compounds such as the indole-containing analytes in this work. Selectivity of the column was such that a marked separation was apparent between 4-HIAA and 5-HIAA-^13^C_6_ and even between PSI and PSI-d10 [34], potentially mitigating the performance of the internal standards. However, it was found that the Waters HSS T3 column provided closer elution of both analyte/IS pairs and also yielded a good separation window between PSI and 4-HIAA (the last eluting metabolite) in which to implement the transition from positive to negative MS ionization. The column also delivered acceptable retention of PSY, the earliest eluting analyte, and good peak shape for all analytes. Therefore, the HSS T3 column was selected for further development and in a 100 mm length to provide greater efficiency and improve resolution between analytes and from matrix interference. Typical aqueous and organic mobile phases (MeCN) modified with 0.1% FA were found to give acceptable retention and peak shape for all analytes without the need for extensive investigation of alternatives.

Extraction of analytes from plasma matrix was performed by protein precipitation. SPE methodologies were investigated but were ultimately unsuccessful due to the polar nature of the analytes and differences in their acid-base characteristics. For example, mixed mode retention of psilocin would be best achieved at low pH (protonating tertiary amine to give cation) whilst 4-HIAA would be best achieved at high pH (deprotonation of carboxylic acid to give anion). Therefore, for the target LLOQs and aim of developing one multi-analyte method, the universal extraction of protein precipitation was preferable over more selective techniques.

Extractions were effected by addition of organic precipitant to the plasma sample in individual tubes or wells of a 96-well plate (polypropylene), followed by mixing, filtration/centrifugation and transfer of the resultant supernatant. The final sample extract needed to be highly aqueous in order to prevent column breakthrough, and peak fronting, upon injection to the chromatography system. This was achieved through evaporation of the organic-rich sample supernatants to dryness, under a stream of nitrogen at 30/40°C, and then reconstituting in water containing 0.1% FA. Once extracted from matrix in organic solvent, analytes did not exhibit instability when handled under standard laboratory conditions. There was also no evidence of analyte instability or interconversion, e.g., hydrolysis of PSY (loss of phosphate) or PSIG (loss of glucuronide) to PSI, during the extraction procedure or dry-down, and supplementation with AA or other antioxidant was not required.

CRO-A employed a Waters Ostro protein precipitation and phospholipid removal plate for extraction from human plasma (method 2). The technique was amenable to higher throughput and should result in cleaner sample extracts (not tested), but no marked benefits in performance or robustness were quantifiable. However, recovery of PSY using the Ostro plate was found to be markedly and unacceptably low in comparison to the other analytes. This is postulated to be attributable to the phosphate group in the molecule interacting with plate sorbent, in a similar manner to the phosphate group present in phospholipids, and being retained. As a consequence, PSY was omitted from the multi-analyte method and a separate single-analyte method developed instead to provide the lowest achievable LLOQ. This method did not employ the Ostro plate in sample extraction and a shorter 50 mm column was used in the chromatographic separation in order to deliver a reduced runtime and increased throughput.

Representative chromatograms obtained for the determination of PSY in blank plasma and plasma fortified with analyte at the LLOQ concentration, employing the final method developed (method 1), are shown in Figure 2. Similarly, representative chromatograms obtained for the determination of PSI, PSIG and 4-HIAA, using the final method developed (method 2), are shown in Figure 3. In chromatograms obtained for PSI, an additional major peak was evident which eluted shortly after the analyte peak itself. This major peak was also present in blank plasma samples and is believed to be attributable to tryptophan [14]. The amino acid is acceptably baseline-resolved from PSI, and does not interfere with its determination. Interestingly, during transfer and establishment of the method at the second vendor, an alternative product ion was selected for detection of PSI (m/z 58 rather than 160) and the tryptophan interference was not observed.

**Figure 2.**
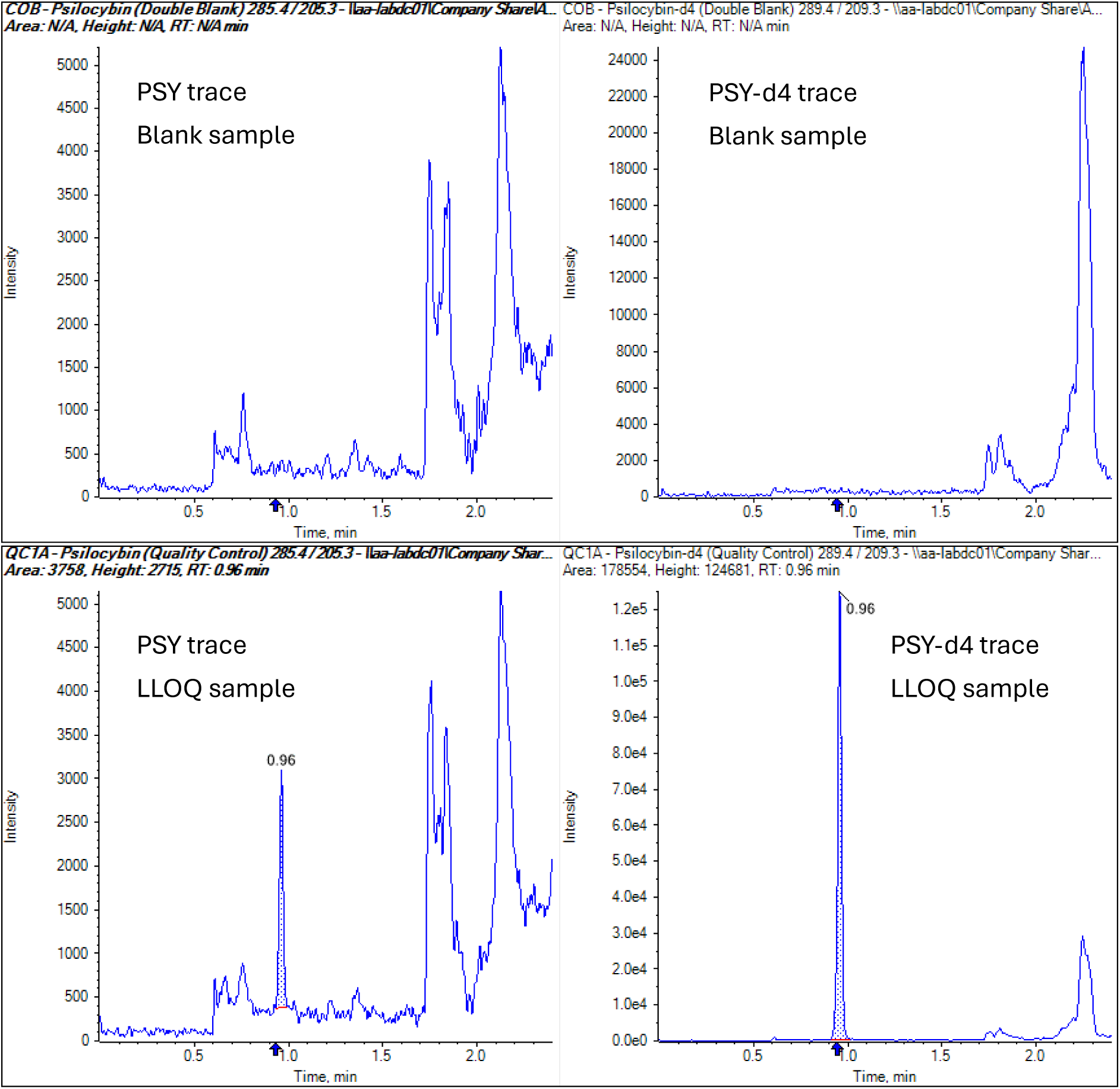
Representative LC-MS/MS chromatograms obtained for PSY and its internal standard, following extraction of blank plasma and blank plasma fortified with analyte at the LLOQ concentration (data generated under method 1).

**Figure 3.**
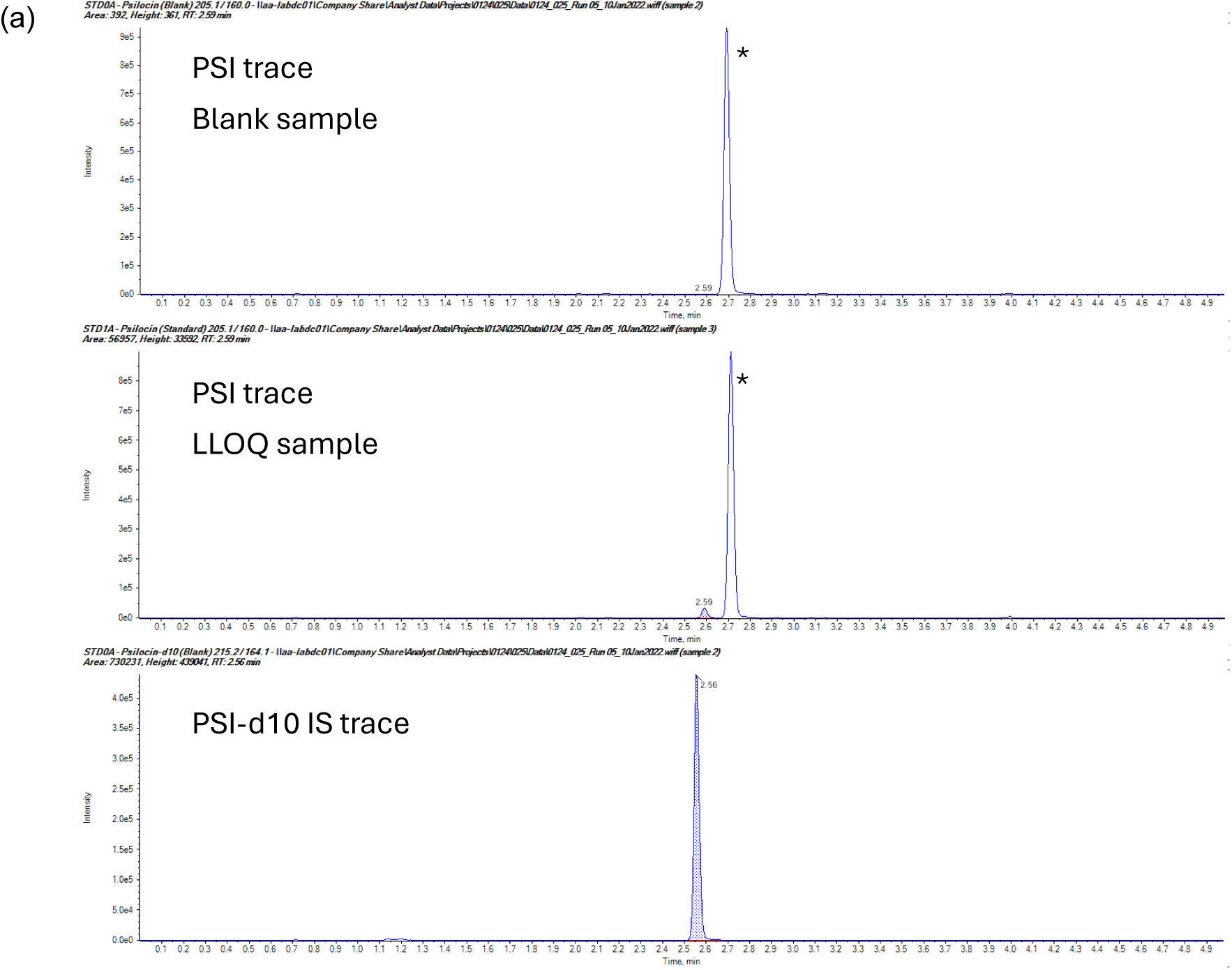
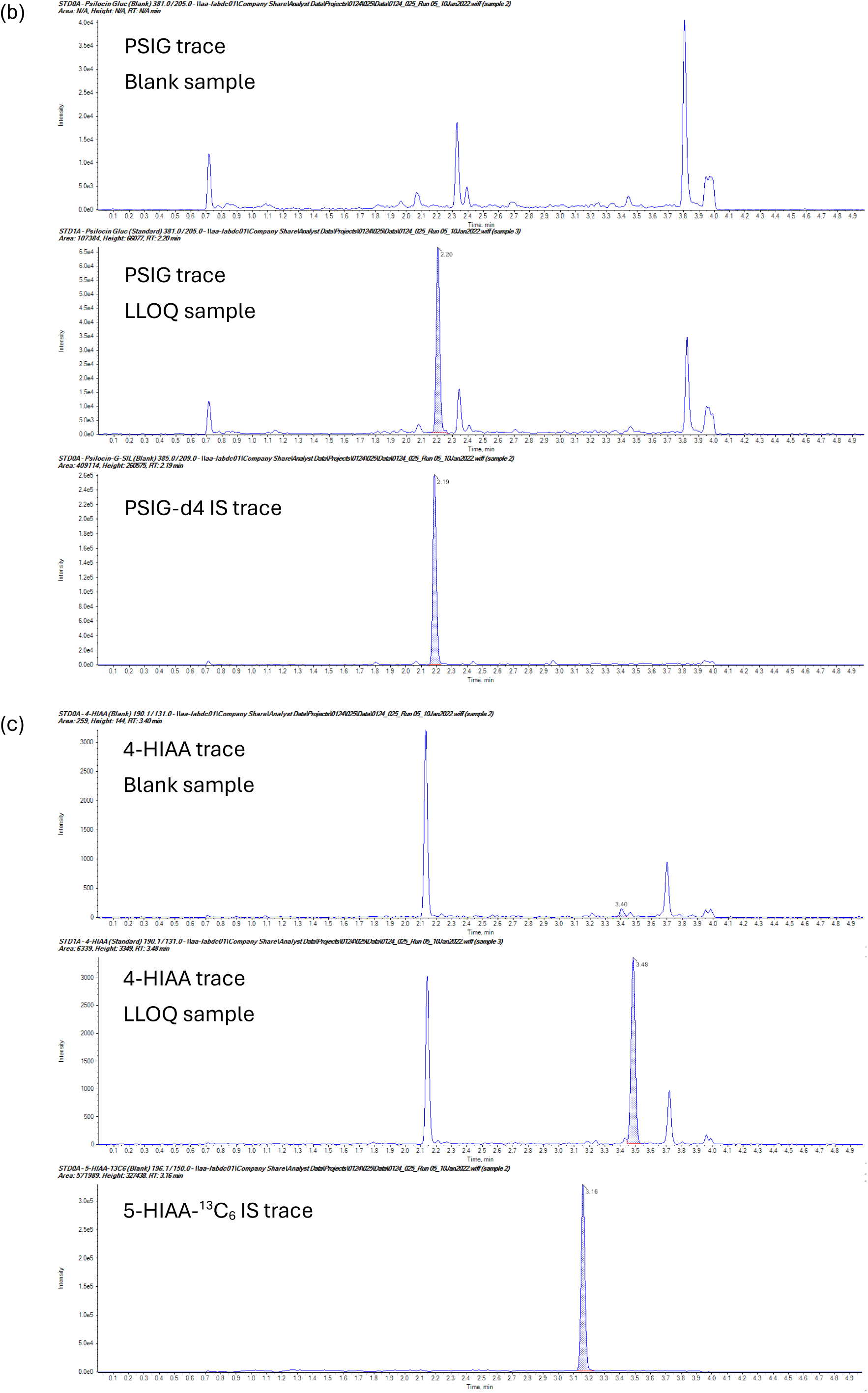
Representative LC-MS/MS chromatograms obtained for (a) PSI, (b) PSIG and (c) 4-HIAA, and their respective internal standards, following extraction of blank plasma and blank plasma fortified with analytes at LLOQ concentrations (data generated under method 2). Major peak (*) present in PSI traces is attributed to endogenous tryptophan.

Robust responses were observed for all analytes, at the selected LLOQ concentrations, and for 4-HIAA in particular. However, as potentially-interfering background peaks were evident in the blank plasma samples tested, which varied from lot to lot, lowering the LLOQ concentration to be validated was discounted.

In agreement with literature, instability of PSI, and to a lesser extent 4-HIAA, in human plasma was evident but the protective effectiveness of AA was unclear; some assessments indicated that PSI stability was decreased in its presence. In practice, it was found that stability could be satisfactorily maintained by keeping samples on ice, avoiding excessive light exposure and minimizing the thawed duration. The absence of a need for adding a stabilising agent is beneficial from the perspective of the clinic where the sample collection protocol can be as straightforward and robust as possible.

Due to unavailability of a stable-labelled version of 4-HIAA at the start of the development program, ^13^C_6_ and d6 versions of the positional isomer 5-HIAA were employed instead. These isomers displayed a slight chromatographic separation between the 2 isomers but performed well as internal standards. However, if a drift in analyte response occurred across a run, on occasion this response was not completely compensated for by the internal standard response and led to an unacceptable higher/lower bias in cal/QC samples at the beginning versus the end of the run. Contract synthesis of a ^13^C_2_,d2-labelled version of 4-HIAA was undertaken and, once available, the material was incorporated into method 3 at CRO-B.

The method for the determination of PSI in human urine was developed from the basis of the plasma assay. Chromatography was modified to be appropriate for the single analyte and deliver a shorter runtime, and sample processing was amended to simple dilution. In this instance, 25mM AA was included in the diluent to protect against potential oxidative losses. Non-specific adsorption to container surfaces, a known potential complication for urine assays, was assessed according to the procedure outlined by Ji et al [35] in which an aliquot of urine spiked with PSI at the mid-QC level was analysed following successive transfers between polypropylene containers; no decrease in response was evident. As PSIG is expected to be present in incurred urine samples at higher concentrations than PSI [6,7], the potential for back-conversion in spiked-samples was assessed and found to be insignificant (<1%) following incubation at ambient temperature for 3 h.

### 3.2 Method validation

#### 3.2.1 Calibration range, accuracy and precision

The concentration ranges employed for analytes and methods, as detailed in Table 1, were validated successfully, applying acceptance criteria as per ICH M10 guidelines. The relationship between nominal analyte concentration and response was found to be linear for PSY with a weighting of 1/x^2^ in method 1. A quadratic regression, with 1/x^2^ weighting, was applied for PSI, PSIG and 4-HIAA in method 2 on the Sciex API6500 instrument. However, a linear regression, with a 1/x weighting, was selected for these analytes in method 3 (and for PSI in human urine in method 4) on the Waters Xevo TQ-XS instrument.

The inter-assay accuracy and precision data for each method are summarized in Table 4. Results are shown for 4-HIAA from validation of method 3 in which both the labelled 5-hydroxy and 4-hydroxy positional isomers were employed as internal standards. Whilst not reflected in the inter-assay results presented, use of 4-HIAA-^13^C_2_,d2 was found to reduce variability within a run; e.g., CV ranged 1.9 to 14.6% (average of 6.0%) and 0.6 to 7.2% (average of 2.4%) employing 5-HIAA-d6 and 4-HIAA-^13^C_2_,d2 respectively. Additionally, an increase in batch acceptance was observed with less failures due to response drift across a run.

#### 3.2.2 Selectivity and specificity

All individual lots of blank plasma tested demonstrated peak area responses of PSY, PSI, PSIG and 4-HIAA, and internal standards, that met acceptance criteria as per guidelines; namely, analyte response < 20% of that at the LLOQ and IS response < 5% of that observed in the LLOQ sample. Haemolysed and lipemic matrix were also free of interference.

Potential interference arising between PSI, PSIG and 4-HIAA in multi-analyte methods 2 and 3 were assessed by individually-spiking each at the ULOQ concentration into a single lot of matrix. The peak area responses of non-spiked analytes were low and, where observed, were approximately 20% of the relevant LLOQ or lower. Psilocin exhibited a response approximately equivalent to the LLOQ in samples individually-spiked with PSIG. It is feasible that the presence of PSI was due to a low level impurity in the reference standard solid material. If not, and the presence arises from deconjugation of PSIG during processing of the sample, the extent of interference is low and would only be impactful in use of the method should a study sample contain a markedly greater concentration of PSIG than PSI (in the order of 100-fold).

Specificity of the methods was demonstrated with respect to a range of potential concomitant medications by experimental (e.g., fluoxetine, sertraline and escitalopram) or *in silico* methods (venlafaxine and its O-desmethyl metabolite) (data not shown).

Lastly, no interfering response was observed for PSI or its IS in 6 individual lots of human urine.

#### 3.2.3 Matrix effect/recovery

The mean recovery of PSY and its IS, from a single source of human plasma in method 1, ranged from 78.4 to 84.4% and from 83.6 to 89.6%, respectively, for the low, mid and high QC concentrations tested. Recoveries determined for PSI, PSIG and 4-HIAA, from human plasma in method 2, were 50.7 to 63.8%, 45.5 to 56.6%, and 65.0 to 72.1%, respectively. The values are slightly low but this may be due in part to non-recovery of the total extract volume through the Ostro extraction plate employed in this method, and recoveries for each IS were broadly consistent at 68.3 to 73.2%, 54.2 to 61.4%, and 73.5 to 78.5%, respectively.

The mean matrix factor determined in methods 1 and 2, for each analyte and respective internal standard, in each source of matrix (n = 6) was generally close to unity at both low and high QC concentrations; i.e., matrix components had no significant impact on MS ionization efficiency. Individual values were between 1.09 to 1.16 for PSY in method 1, and 1.00 to 1.26, 1.21 to 1.61, and 1.05 to 1.23 for PSI, PSIG and 4-HIAA, respectively, in method 2. It follows that the IS-normalised matrix factor for each analyte was also close to unity and the precision across all sources of each matrix was less than 15%. The IS-normalised matrix factor was consistent in plasma sources that were haemolysed (≥2% haemolysis) or lipemic.

The overall matrix effect determined by spiking individual sources of human plasma (n ≥ 6) with PSI, PSIG and 4-HIAA, at low and high QC concentrations in method 3, met regulatory acceptance criteria for both accuracy and precision. Accuracy across the 6 lots of matrix, and haemolysed and lipemic sources, for PSI, PSIG and 4-HIAA were 88.0 to 109.4%, 86.7 to 106.9% and 96.3 to 110.6%, respectively; precision was 0.85 to 4.6%, 1.6 to 4.1% and 2.0 to 11.2%, respectively.

The matrix effect accuracy and precision for determination of PSI in individual sources of human urine (n ≥ 6), at low and high QC concentrations in method 4, also met regulatory acceptance criteria with values of 88.7 to 97.5% and 1.2 to 6.2%, respectively.

Overall, whether assessment was performed through determination of recovery and matrix factor separately, or through the overall matrix effect, results demonstrated that the methods were not impacted by variance in the source of matrix.

#### 3.2.4 Stability

The results from stability assessments on the analytes, under various conditions of sample handling and storage, are presented in Table 5.

All analytes demonstrated acceptable stability in whole blood, prior to plasma preparation, if kept in wet ice.

Psilocybin was found to be relatively stable in human plasma and if handled/thawed on wet ice showed no marked decrease in determined concentration following storage for at least 4h or following 6 freeze/thaw (F/T) cycles. Stability of PSY was also shown in samples stored frozen for at least 1 or 6 months at -20 or -80°C, respectively.

The intrinsic stability of PSI, PSIG and 4-HIAA in human plasma is best demonstrated from results of tests performed in thawed matrix at room temperature; the concentrations of PSIG and 4-HIAA determined were acceptable following storage of up to 20h, whilst those of PSI had decreased to ∼74% after 6h and ∼33% after 20h. Correspondingly, PSIG and 4-HIAA were found to be acceptably stable in human plasma following at least 5 F/T cycles (thawed on wet ice) and stored frozen for at least 2 or 9 months at -20 or -80°C, respectively. PSI showed low, yet acceptable, bias following 5 F/T cycles and frozen storage at -20°C for 1 month, but an unacceptably low accuracy (∼19 or ∼84%) following frozen storage at -20°C for 2 months or -80°C for 9 months, respectively; acceptable stability was, however, demonstrated following frozen storage at -80°C for 6 months

Acceptable stability of PSI in human urine was also demonstrated under conditions reflective of sample collection and storage. For example, an acceptable bias of ∼88% was observed following storage in a collection vessel at 4°C for 27h.

Additional stability assessments (data not shown) under the conditions discussed were also conducted on dilution QCs, in the event that study sample results were above the calibration range. A general trend of decreased stability of PSI at higher concentrations was evident. For example, dilution QC samples containing PSI at 4000 ng/mL showed an accuracy of ∼54% following storage on wet ice for 8h, compared to ∼89% for low/high QC samples. As determined concentrations in study samples were within the calibration range, this finding was not investigated further.

Lastly, extract stability was assessed and demonstrated acceptable reinjection reproducibility (data not shown) following storage of extracts at 10°C for at least 143h for PSY (in method 1) and 109h for PSI/PSIG/4-HIAA (in method 2).

#### 3.2.5 Method application and ISR

Successful application of the methods has been demonstrated, at both vendors, in support of 6 clinical studies. Owing to instability of the analytes, robust adherence to sample collection, handling and storage processes is required and high operational performance of instrumentation is needed in analysis of the samples.

Representative plasma concentration data from a cohort of 6 healthy participants receiving a single oral dose of 25 mg COMP360, under fasting conditions, are presented in Figure 4.

**Figure 4.**
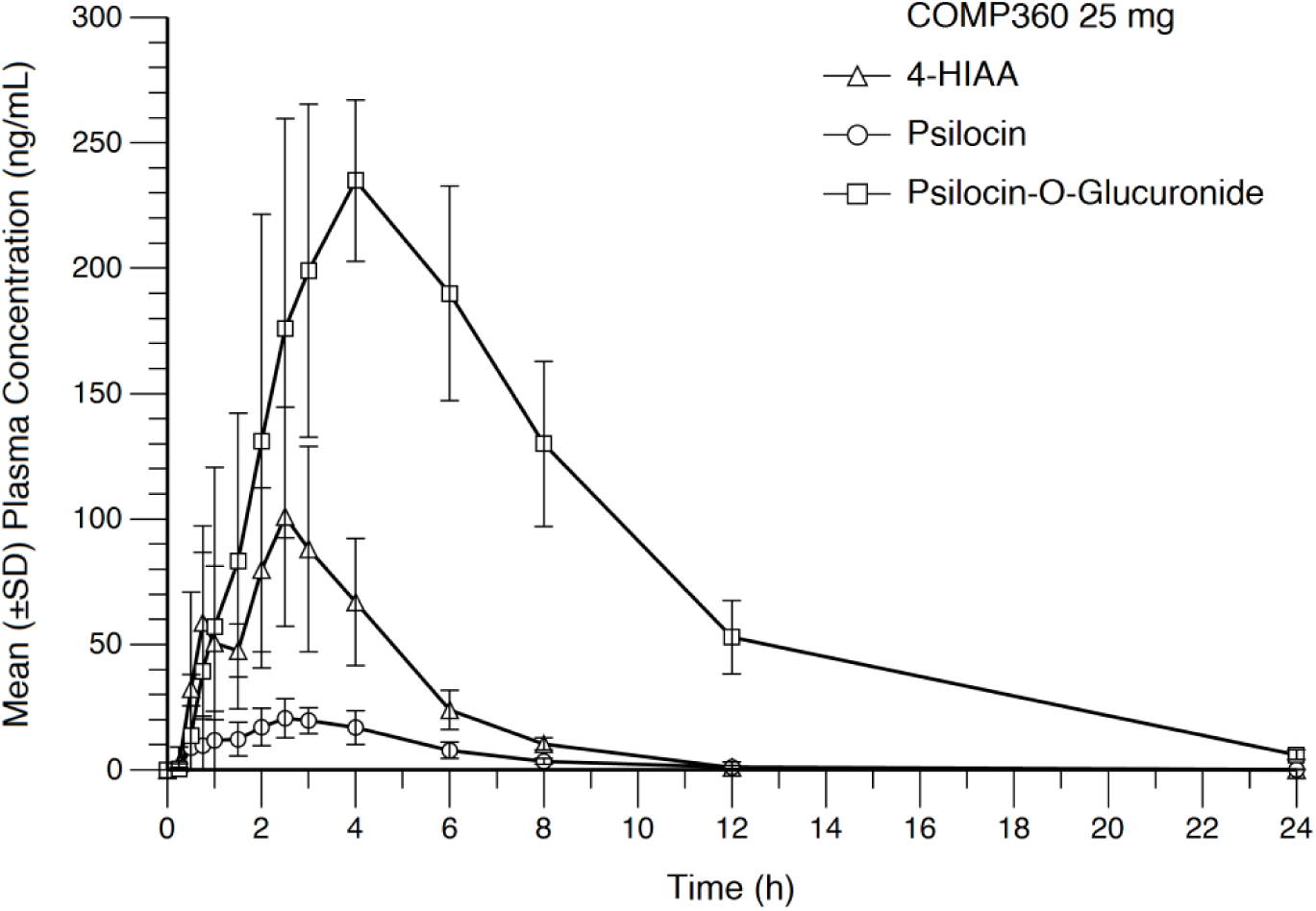
Concentration-time plots determined for PSI, PSIG and 4-HIAA in plasma from healthy volunteers administered an oral dose of 25 mg COMP360

The validated method LLOQs allowed for concentrations of PSI (the active species) to be determined out to 12h, and for concentrations of 4-HIAA and PSIG (major metabolites) out to (at least) 8 and 24h, respectively. The validated method ULOQs were also found to be appropriate with no samples (from the 25 mg dose) requiring reanalysis with dilution for any analyte.

PSY was unquantifiable (< 0.500 ng/mL) in all samples from participants receiving the proposed therapeutic dose of 25 mg or the supratherapeutic dose of 50 mg. Operationally, this aligned with the methods developed and validated in that it was only necessary to analyse samples using the multi-analyte method to determine concentrations of PSI, PSIG and 4-HIAA in subsequent trials.

The concentrations determined for psilocin [7,13,17,18,36] and 4-HIAA [13,36], and the unquantifiable results for PSY [17], were consistent with other published studies following oral administration of PSY at doses of 25 mg or similar weight-based doses. However, the concentrations determined for PSIG were noticeably greater than those reported elsewhere [13,36]. This disagreement is believed to be attributable to differences in the procedure through which PSIG concentrations have been generated. In this work, concentrations in study samples have been determined directly in comparison to known concentrations of PSIG spiked into calibration (and QC) samples. In the other work, PSIG has been quantified indirectly by determining PSI concentrations in samples before and after enzyme hydrolysis and comparing the results. In part, the non-equivalence is believed to be attributable to the manner in which the concentrations of PSIG are represented; in this work, concentration simply refers to the mass of PSIG per unit volume whereas, in other work, reflecting how the value was determined, it is expressed as concentration equivalents of PSI (i.e., not normalised for difference in molecular weight of PSI and PSIG). Further, as alluded to earlier, indirect quantification of PSIG through enzyme hydrolysis may also result in underestimation of the actual concentration due to incomplete hydrolysis and/or degradation of PSI under the incubation conditions required to effect deconjugation.

A summary of the results from the ISR analysis performed, across clinical studies, methods and matrices, are presented in Table 6 and demonstrate acceptance with regulatory guidance (at least 2/3 of samples within acceptance limit).

**Table 6.** Summary of ISR results determined in clinical studies.

| Study | Matrix | Method | Percentage of acceptable ISR results* (number of samples) |  |  |  |
|---|---|---|---|---|---|---|
|  |  |  | PSY | PSI | PSIG | 4-HIAA |
| Healthy volunteers in clinical study 1 | Plasma | 1, 2 | ND | 84.7% (59) | 95.1% (81) | 82.4% (51) |
| Healthy volunteers in clinical study 2 | Plasma | 2 | N/A | 92.9% (42) | 93.8% (48) | 94.9% (39) |
| Healthy volunteers in clinical study 3 | Plasma | 3 | N/A | 71.6% (102) | 90.0% (100) | 86.3% (102) |
| Patient participants in clinical study 4 | Plasma | 2 | N/A | 86.7% (45) | 80.9% (47) | 97.6% (41) |
| Healthy volunteers in clinical study 5 | Urine | 3 | N/A | 100% (14) | N/A | N/A |

\* Result acceptable if percent difference within ±20%
ND Not determined – all study samples were below limit of quantification
N/A Not applicable

#### 3.2.6 Human plasma method cross-validation

Due to human plasma concentration data being generated across studies using different fully validated methods (method 2 and 3), and this data being integrated to support regulatory decision making, a cross-validation of the methods was conducted. The evaluation employed spiked QC samples and incurred samples from a clinical study.

The cross-validation QC samples were prepared at one vendor and analysed at both, and the determined concentrations of each analyte, at both laboratories, met standard acceptance criteria for intra-assay batch accuracy and precision. Across all analytes, accuracy ranged from 102.3 to 111.2% and 94.4 to 108.7%, and precision from 1.7 to 4.5% and 1.3 to 8.8%, at vendor laboratories CRO-A and CRO-B, respectively.

A plot of the %deviation obtained for each cross-validation QC sample and the incurred samples is presented in Figure 5. Excepting 2 outlier results (one in the QC sample set and the other in the incurred sample set), data show excellent agreement in concentrations between laboratories. Evaluation of the sample sets in the same manner as ISR analysis would be conducted within a study [37], it was found that 94.4% of spiked QC samples (for all analytes) and 92.9, 93.3 and 96.4% of incurred samples (for PSI, PSIG and 4-HIAA, respectively) deviated by less than ±20%.

**Figure 5.**
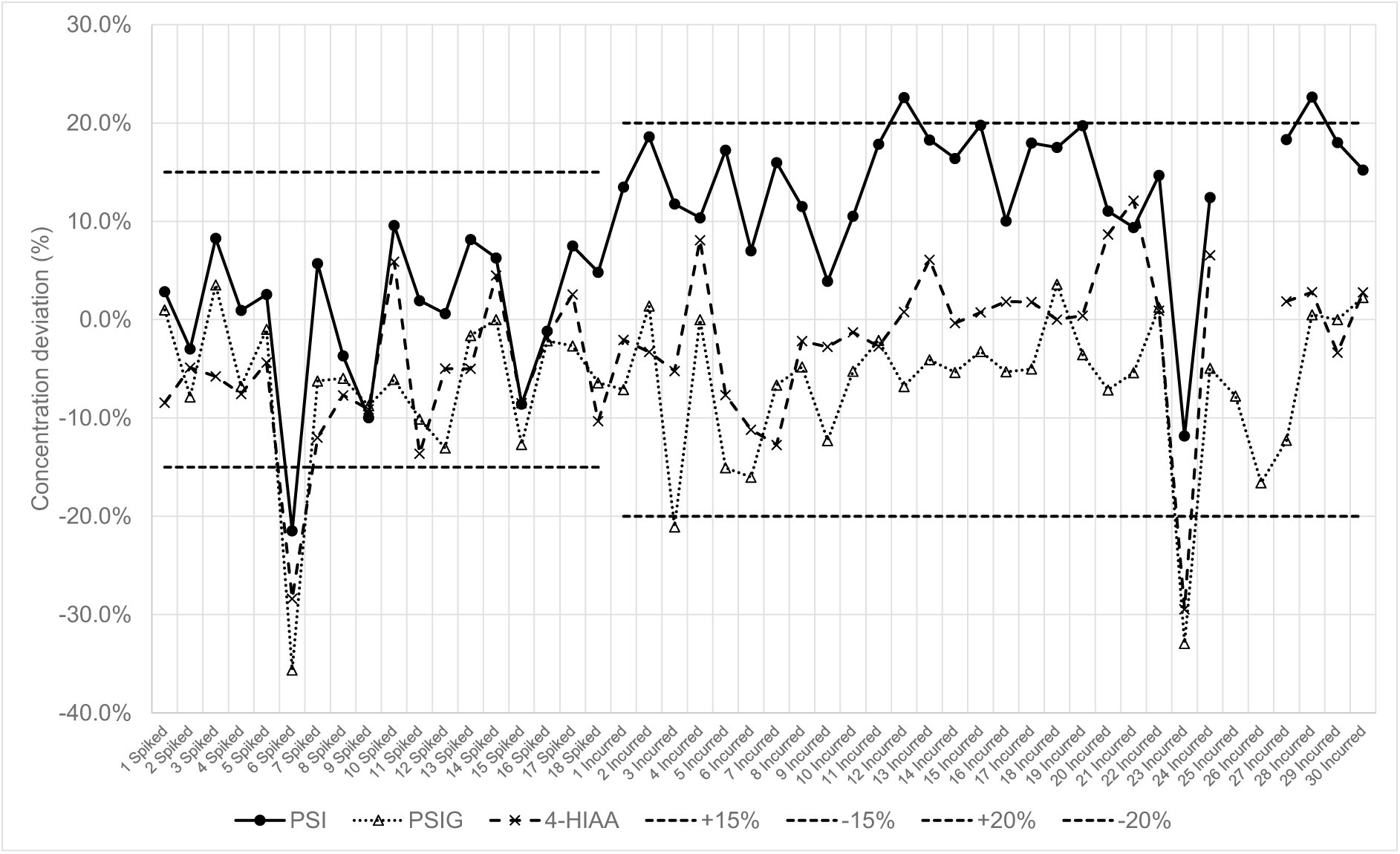
Plot of %deviation of determined concentration, comparing CRO-A and CRO-B, in each cross-validation sample (spiked QC and incurred) analysed

Regulatory guidance on cross-validation (ICH M10) has moved away from specified acceptance criteria *per se* and placed emphasis on performing an assessment of agreement between methods and of impact on data sets. This has been reflected in several articles released subsequently [see e.g., 38-40]. Bland-Altman plots of %deviation of determined concentration, comparing CRO-A and CRO-B, against the mean concentration in each cross-validation sample analysed is presented in Figure 6. The mean %deviation for PSI, PSIG and 4-HIAA in incurred samples was assessed to be 13.9%, -6.7% and -1.0% (CRO-B vs CRO-A), respectively, and there was no evidence of the %deviation being concentration-dependent for any analyte.

**Figure 6.**
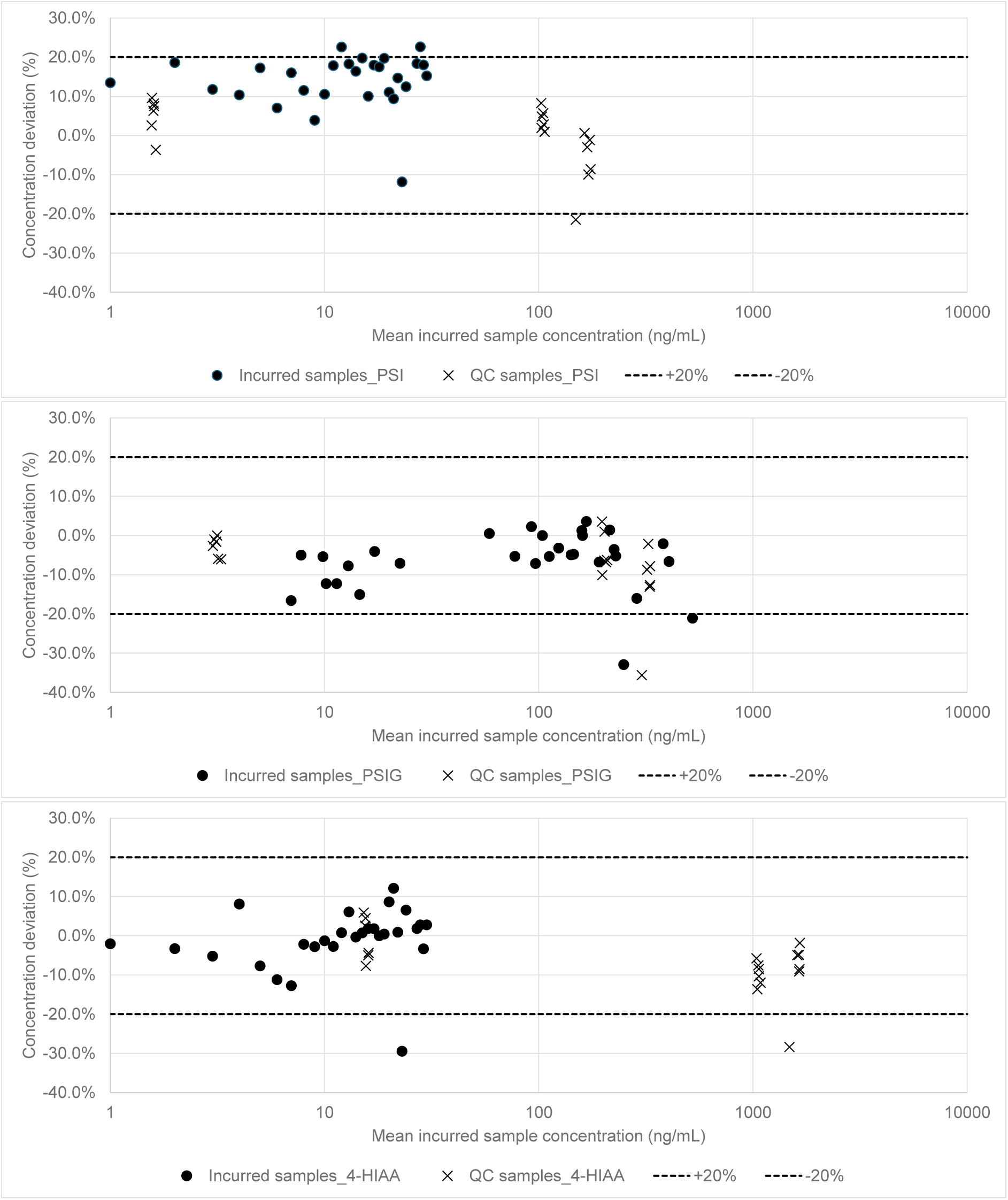
Bland-Altman plot of %deviation of determined concentration, comparing CRO-A and CRO-B, against the mean concentration in each cross-validation sample analysed

## 4. Conclusion

Bioanalytical methods, employing LC-MS/MS, have been developed for the determination of PSY, PSI, PSIG and 4-HIAA in human plasma, and PSI in human urine across 2 vendor bioanalytical laboratories. The methods have been fully validated to satisfy the requirements of ICH M10 guidance and acceptable stability results for all analytes have been demonstrated under the conditions of sample handling and storage. The methods have been successfully employed in analysis of test samples from 6 clinical studies to determine pharmacokinetic concentrations following administration of COMP360, a form of synthetic PSY. The reproducibility of study concentrations has been demonstrated through successful ISR analysis. Comparative performance of the multi-analyte method for PSI/PSIG/4-HIAA, across vendor laboratories, has been shown through the conduct of a cross-validation study employing spiked-QC and incurred samples. These validation and sample analysis activities have been pivotal in the development of COMP360.

## CRediT authorship contribution statement

**S. Silvester:** Conceptualization, Methodology, Validation, Formal analysis, Writing - Original Draft, Visualization, Supervision, Project administration. **L. Wray:** Writing - Review C Editing, Supervision, Project administration. **H. Dooner:** Writing - Review C Editing, Visualization, Supervision. **G. Gilmour:** Writing - Review C Editing. **A. Patel:** Writing - Review C Editing.

## Acknowledgements

The authors wish to acknowledge scientists at Alderley Analytical (Synexa Life Sciences, UK) and Quotient Sciences Alnwick (Arcinova, UK) who performed the method development, validation activities and sample analysis described within this article.

